# Projection of Cortical Beta Band Oscillations to a Motor Neuron Pool Across the Full Range of Recruitment

**DOI:** 10.1101/2025.03.18.644010

**Authors:** E Abbagnano, A Pascual-Valdunciel, B Zicher, J Ibáñez, D Farina

## Abstract

Cortical beta band oscillations (13–30 Hz) are associated with sensorimotor control, but their precise role remains unclear. Evidence suggests that for low-threshold motor neurons, these oscillations are conveyed to muscles via the fastest corticospinal fibers. However, their transmission to motor neurons of different sizes may vary due to differences in the relative strength of corticospinal and reticulospinal projections across the motor neuron pool. Consequently, it remains uncertain whether corticospinal beta transmission follows similar pathways and maintains consistent strength across the entire motor neuron pool. To investigate this, we examined beta activity in motor neurons innervating the tibialis anterior muscle across the full range of recruitment thresholds in a study involving 12 participants of both sexes. We characterized beta activity at both the cortical and motor unit levels while participants performed contractions from mild to submaximal levels. Corticomuscular coherence remained unchanged across contraction forces after normalizing for the net motor unit spike rate, suggesting that beta oscillations are transmitted with uniform strength to motor neurons, regardless of size. To further explore beta transmission, we estimated corticospinal delays using the cumulant density function, identifying peak correlations between cortical and muscular activity. Once compensated for variable peripheral axonal propagation delay across motor neurons, the corticospinal delay remained stable, and its value (approximately 14 *ms*) indicated projections through the fastest corticospinal fibers for all motor neurons. These findings demonstrate that corticospinal beta band transmission is determined by the fastest pathway connecting in the corticospinal tract, projecting uniformly across the entire motor neuron pool.

## Introduction

Beta oscillations (13–30 Hz) are a prominent neural rhythm in the sensorimotor cortex and are associated with steady motor states and sensorimotor integration (Baker, 2007; Kilavik et al., 2013). These oscillations are transmitted to muscles during sustained motor tasks, as shown by corticomuscular coherence (CMC) analysis (Baker, 2007; Conway et al., 1995; Witham et al., 2011). This coupling between the cortex and muscles in the beta band indicates phase synchronization between cortical and muscular activity, evident especially during isometric contractions (Baker et al., 1997; Echeverria-Altuna et al., 2022; Salenius et al., 1997). Recent studies have shown that beta oscillations are transient rather than sustained neural signals, appearing as phase-coupled brief bursts at both cortical and muscular levels (Bonaiuto et al., 2021; Echeverria-Altuna et al., 2022; Little et al., 2019).

While it has been proposed that peripheral beta activity does not directly influence volitional force modulation, further investigation into its broader role in motor control is needed (Zicher et al., 2024, 2023). Previous work has shown that these oscillations exhibit abnormal corticomuscular coupling in neurological injuries associated with motor impairments (Engel and Fries, 2010), such as Parkinson’s disease, stroke, and spinal cord injuries, reflecting disrupted communication between the cortex and muscles (Gourab and Schmit, 2010; Von Carlowitz-Ghori et al., 2014; Zokaei et al., 2021). Thus, while the exact function of peripherally transmitted beta oscillations remains uncertain, their transmission is likely crucial for motor control (Baker et al., 1999).

Cortical beta oscillations are transmitted through the corticospinal tract, with signals propagating along descending axons and summing at the soma of lower motor neurons (MN) (Baker et al., 2003). The conduction delays in this tract suggest that, at least for low-threshold MNs, beta transmision is determined by the fastest pathways in the corticospinal tract, as these contribute most significantly to the synchronization observed between cortical and peripheral beta rhythms (Ibáñez et al., 2021). However, while it is known that the input to a MN pool is largely shared across all MNs at *low* frequencies (Farina et al., 2014), which are associated to force control, it remains unclear whether beta band inputs follow the same distribution and pathways across different MN sizes.

Lower MNs within a pool vary in size and electrophysiological properties, following the size principle during voluntary activation (Heckman and Enoka, 2004; Henneman, 1957). While beta oscillations have been shown to be transmitted to peripheral muscles via low-threshold MNs, which are primarily involved in fine motor control, their transmission to high-threshold MNs engaged in strong contractions remains unclear. These units appear to be more influenced by the reticulospinal than the corticospinal tract (Baker, 2011; Glover and Baker, 2020). Accordingly, previous studies have reported a decrease in CMC values within the beta band as contraction force increases (Cremoux et al., 2017; Dal Maso et al., 2017; Ushiyama et al., 2012). This decline may indicate either a reduction in the common beta band input at higher contraction forces or a different projection of beta on small and large MNs. However, standard electromyography (EMG) and CMC analyses face limitations due to amplitude cancellation at high forces (Day and Hulliger, 2001; Farina et al., 2008; Keenan et al., 2006). These limits can be partially overcome by decomposing the EMG signal into the discharge activations of individual motor units (MUs) (Farina and Holobar, 2016).

In this study, we characterized the projection of beta band oscillations to the pool of spinal MNs innervating the Tibialis Anterior (TA) muscle. We first demonstrated that we could identify the activity of MNs across a large range of recruitment thresholds. Despite the differences in thresholds, our results showed that beta oscillations were transmitted to all MNs using the fastest corticospinal fibers and with an approximately uniform strength, irrespective of MN size. These results provide novel insights into the transmission of beta oscillations to spinal MNs, which could ultimately contribute to elucidating the role of beta oscillations in motor control.

## Methods

### Experimental Data Acquisition

#### Subjects

Twelve healthy participants (ages: 28 ± 4.63 years, 10 males and 2 females) with no known history of neurological or musculoskeletal disorders were recruited for this study. While this sample size is comparable to previous studies (Bräcklein et al., 2022; Ushiyama et al., 2012; Zicher et al., 2024), the analysis method we employed provided a more precise assessment of corticospinal transmission by directly analysing individual MUs rather than relying solely on global interference EMG (see below description of methods). This granular analysis reduces variability and improves sensitivity in detecting differences, leading to highly consistent and reliable findings, as demonstrated in the Results section. All participants provided written informed consent prior to their inclusion. The study was conducted in accordance with the principles outlined in the Declaration of Helsinki and received approval from the Imperial College London Ethics Committee (reference number 18IC4685).

#### Experimental Paradigm

To investigate the transmission of beta band activity across the MN pool, it is essential to study a broad range of MNs, which vary in size and recruitment thresholds. Larger MNs have higher thresholds than smaller MNs, thus require larger excitatory input to initiate firing (Henneman, 1957). Therefore, the experimental design involved subjects performing isometric contractions at progressively increasing forces, determining the recruitment of MNs across a large range of forces (Henneman, 1957). Isometric contractions were selected because they enhance beta band power both cortically and peripherally (Baker, 2007; Engel and Fries, 2010) while also facilitating the identification of individual motor unit firings at a steady rate. Cortical activity was measured using electroencephalography (EEG), while high-density surface electromyography (HD-sEMG) was used to record signals from the TA muscle. The experimental paradigm is visualized in Figure 1.

**Figure 1.**
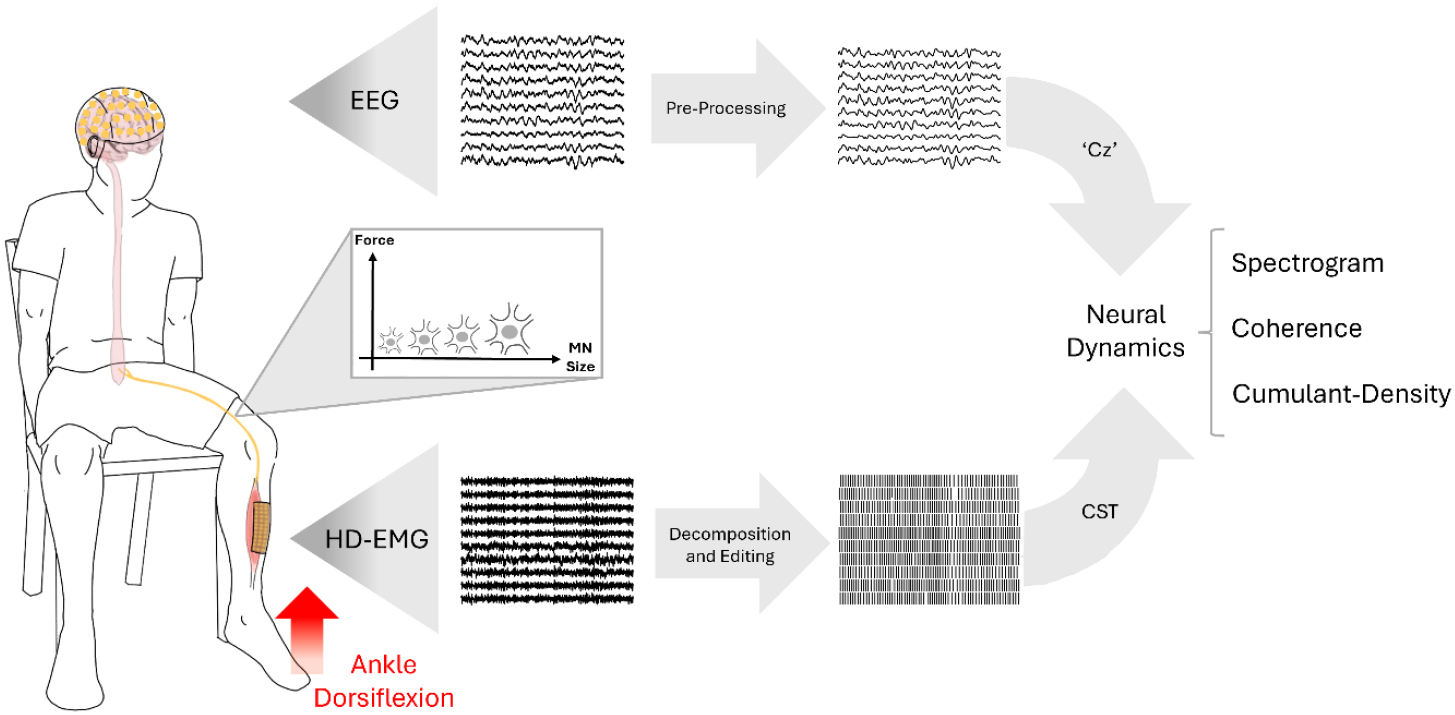
Overview of the experimental paradigm. Subjects performed multiple isometric dorsiflexion contractions using the tibialis anterior muscle to investigate the transmission of cortical beta band oscillations to the muscle. Contractions were performed at 5%, 10%, 20%, 30%, 50%, and 70% of the maximum voluntary contraction, progressively recruiting larger motor neurons in accordance with Henneman’s size principle. During each contraction, EEG and high-density EMG (HD-EMG) were recorded simultaneously to examine transmission across different motor neuron sizes. EEG signals were filtered offline, artifacts were removed, and the ‘Cz’ channel was selected for further analysis. HD-EMG was decomposed into individual motor unit spike trains, from which the cumulative spike train (CST) was computed. Transmission of cortical beta band oscillations between cortex and muscle was then analysed using spectrograms, coherence, and cumulant density.

#### Experimental Design

Participants were seated comfortably in a chair with their knee flexed at approximately 75°, and their leg securely fixed to an ankle dynamometer using straps. The foot was positioned on a pedal inclined at 30° in the plantarflexion direction (where 0° corresponds to the foot being perpendicular to the shank). At the start of the session, participants performed a single maximum voluntary contraction (MVC) of the ankle with a dorsiflexion to determine their individual MVC, which served as a reference for the subsequent tasks. Visual feedback was provided throughout the experiment, with a target displayed on a screen representing the required force level, and a real-time trace showing the force produced by the participant.

Each participant performed a total of 16 isometric dorsiflexion contractions with the TA. These included a linear ramp phase where force increased from 0% to the target MVC level at a rate of 5% MVC per second, followed by a plateau phase at the target level, and finally a ramp-down phase back to 0% MVC at the same rate (trapezoidal force profile). Six target MVC levels were used: 5%, 10%, 20%, 30%, 50%, and 70%. For the lower targets (5%, 10%, 20%, and 30%), the plateau phase lasted 60 seconds. To minimize fatigue, contractions at 50% and 70% MVC were subdivided into multiple repetitions: five repetitions of 15 seconds for 50% MVC and seven repetitions of 10 seconds for 70% MVC. The duration of these plateaus was chosen based on previous studies showing successful decomposition for segments longer than 5 seconds at sub-maximal contraction levels (Avrillon et al., 2024b; Del Vecchio et al., 2019). The sequence of these contractions was randomized across participants. There was a 2-minute rest period in between each contraction to reduce fatigue.

#### Data Acquisition

HD-sEMG signals were recorded from the TA muscle of the self-reported dominant leg using a 256-electrode grid (10 columns × 26 rows, gold-coated, 1 mm electrode diameter, 4 mm inter-electrode spacing; OT Bioelettronica). The electrode array was positioned over the muscle belly, aligned with the muscle fibers direction. EMG signals were recorded in monopolar derivation, amplified using the Quattrocento Amplifier system (OT Bioelettronica, Torino, Italy), sampled at 2048 Hz and digitally band-pass filtered between 10 and 500 Hz. Force data were collected using a transducer (TF-022, CCT Transducer s.a.s) mounted on the pedal of the ankle dynamometer and digitalized at 2048 Hz using the Quattrocento Amplifier system.

EEG signals were recorded concurrently using 31 active gel-based electrodes placed according to the international 10–20 system (actiCAP, Brain Products GmbH). The FCz channel served as the online reference. EEG signals were amplified with the BrainVision actiCHamp Plus system (Brain Products GmbH), sampled at 1000 Hz, and subsequently resampled to 2048 Hz.

To ensure precise temporal alignment, all recordings were synchronized using a shared digital trigger signal sent to both the Quattrocento Amplifier system and the BrainVision actiCHamp Plus.

### Data Analysis

#### HD-sEMG decomposition and processing

HD-sEMG signals recorded during the experimental tasks were offline decomposed to identify the activity of individual MUs using a convolutive blind source separation algorithm (Negro et al., 2016). To ensure high-quality data, spike trains with a silhouette value (SIL) below 0.9 were automatically discarded (Holobar et al., 2014; Negro et al., 2016). The decomposed spike trains were then manually reviewed to identify and correct errors. Adjustments were made by recalculating the motor unit separation filters and reapplying them to the raw EMG data to refine the spike train estimation, following well-established guidelines (Del Vecchio et al., 2020).

MU properties were extracted using a validated MATLAB script (Avrillon et al., 2024b). Key properties included the average smoothed firing rate and the recruitment threshold, defined as the percentage of MVC at which the MU began firing.

To obtain longer spike trains for analysis at 50% and 70% MVC, individual MUs were tracked across repetitions. Tracking was performed using a validated MATLAB algorithm that leveraged the unique spatial distribution of motor unit action potential (MUAP) waveforms (Avrillon et al., 2024b). Pairing between MUs in different contractions was retained only if their MUAP cross-correlation exceeded 0.9 (Martinez-Valdes et al., 2017). However, regardless of the cross-correlation value, pairs were rejected if recruitment thresholds differed by more than 10% MVC, as this indicated a potential mismatch. After tracking, spike trains from successfully matched MUs across repetitions were merged.

Following these procedures, additional quality control criteria were applied. MUs were excluded if their SIL was below 0.9, their firing duration was less than 50% of the task time, or their instantaneous firing rate had a standard deviation exceeding 20 Hz (Del Vecchio et al., 2020; Negro et al., 2016). Decomposition and manual editing were conducted using the MUedit MATLAB application (Avrillon et al., 2024a). Only MUs meeting all quality criteria were retained for further analysis.

Because of the variability in the decomposition algorithm to define the discharge time, discharges were realigned to the MUAP onset, using the methods described in Ibáñez et al. (2021).

Following decomposition, the individual MU spike train was represented as a binary signal, where a value of 1 indicated a firing event. By summing these binary signals across a pool of decomposed MUs, a Cumulative Spike Train (CST) was obtained, providing an estimate of the neural drive conveyed by MNs to the muscle (Farina and Negro, 2015). For each contraction level, a distinct CST was computed, resulting in six CSTs per participant, one for each contraction level.

To ensure that CSTs from different contraction levels, which may differ in the number and properties of contributing MUs, were comparable when used in the following coherence analyses (see below), it was necessary to standardize the amplification of the common input projected to the MN pool. In fact, the amplification gain of the common input in a CST is determined by the total number of MU firings, which is given by the number of recruited MUs and their firing rates (Farina et al., 2014). To address this issue, CSTs were normalized by ensuring that the total number of firings was matched across all contraction levels for each participant. Specifically, the contraction level with the lowest total number of firings was identified, and MUs were randomly selected from other levels to match this firing count. This random selection was repeated 10 times for each contraction level, generating 10 permutations of CSTs. In most cases, the contraction levels with the lowest firing counts—where no permutations could be performed—were 50% or 70% MVC. However, results indicated that variance remained consistent across contraction levels, suggesting that this normalization did not introduce significant differences in variability of the estimates.

All subsequent analyses were conducted on these CSTs, and the final values reported for each subject and contraction level were computed as the average across the 10 permutations.

#### EEG processing

The preprocessing of EEG signals was performed using the FieldTrip Toolbox (Oostenveld et al., 2011). Signals were first offline re-referenced to the average of the earlobe electrodes recorded during the experiment. A fourth-order Butterworth bandpass filter was applied to retain frequencies between 0.5 and 45 Hz. Independent Component Analysis was then utilized to identify and remove artifacts caused by eye blinks, oculomotor activity, muscle activity, and other noise sources. To further enhance signal quality, a surface Laplacian filter was applied to eliminate common inputs across electrodes using the CSD Toolbox (Kayser and Tenke, 2006).

For subsequent analysis, the ‘Cz’ channel was selected as it typically exhibits the strongest CMC in the beta band during TA activity (Ibáñez et al., 2021; Zicher et al., 2023). In the connectivity analysis, the ‘Cz’ signal underwent additional correction to account for the phase shift inherent to the pyramidal tract (Baker et al., 2003). This adjustment was performed by computing and inverting the first derivative of the processed ‘Cz’ signal, a method shown to improve the accuracy of delay estimation between cortical and muscular signals and remove confounding factors (Ibáñez et al., 2021).

#### Spectral Analysis

The projection of cortical beta band oscillations to the TA was evaluated using connectivity analysis between the processed ‘Cz’ EEG signal and the CSTs derived as previously described. This analysis was performed using the Neurospec 2.11 toolbox for MATLAB (www.neurospec.org, Halliday, 2015). Coherence was estimated using 1-s signal segments and multitaper spectral estimation (three tapers). The upper 95% confidence limit for significance was determined as 1 −0.05^1/*(L*−1)^, where *L* is the number of segments used in the analysis (Rosenberg et al., 1989). To ensure uniform coherence significance thresholds across conditions, all the contraction plateau phases were cropped to the same length. This resulted in 49 aligned segments per subject and contraction level, establishing a significance threshold of 0.041.

Intramuscular coherence (IMC) was also computed to investigate its association with CMC. For IMC estimation, the MU pool was divided into two randomly selected subsets of equal size. This process was repeated over 50 iterations, with a different configuration of subsets for each iteration. The IMC was calculated as the average coherence between pairs of subsets across all iterations.

Corticomuscular transmission was further characterized in the time domain using the cumulant density function, which estimates the transmission delay at which significant correlation between cortical and muscular activity peaks (Halliday et al., 1995). Similar to coherence, confidence limits for significant correlation peaks were defined as 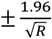, where *R* represents the number of data points. To investigate corticomuscular transmission delays originating from the brain, the analysis targeted the time point of the highest peak identified at positive lags. If this peak was not statistically significant, no transmission delay was assigned for that subject at the given contraction level.

Power spectral density (PSD) was calculated using Welch’s method in MATLAB with the function *pwelch* (4-second window, 75% overlap). This analysis provided insight into the spectral properties of the cortical and muscular signals.

#### Beta Bursting Activity

Beta bursting activity was analysed to assess whether its features varied with force levels or the size of MNs transmitting the common input. Beta bursts in the EEG signal and the CST were identified by applying a thresholding method. Both signals were bandpass filtered (13–30 Hz) using a fourth-order Butterworth filter, and an initial threshold, based on the median envelope value, was incrementally adjusted for both signals (0 to 6 times the median in 0.25 steps). Their envelopes were segmented into 1-s intervals, and the Pearson correlation coefficient was computed for the power of the envelope and the percentage of the envelope exceeding the threshold. This process was repeated for every given threshold and then averaged across blocks. The optimal threshold was identified as the one with the highest correlation value. Beta bursts were defined as intervals where the signal envelopes exceeded this threshold (Little et al., 2019; Shin et al., 2017). Burst rate was quantified as the average number of bursts per second, while burst duration was calculated as the average time the envelope remained above the threshold.

Following Bracklein et al. (2022)., we classified periods above the threshold as ON events and the intervals between two consecutive ON events as OFF periods. To investigate peripheral burst occurrences in relation to cortical ON events, we identified cortical bursts for each subject and paired them with peripheral windows of equal duration, adjusted for the estimated average transmission delay across subjects at each contraction level calculated using cumulant density analysis. We then averaged peripheral beta activity during these delayed ON windows and compared it to both OFF periods and non-delayed ON windows. Only significant correlation values were reported across subjects in this analysis.

#### Statistical Analysis

All statistical analyses were conducted using Jamovi software (www.jamovi.org) and custom-written MATLAB scripts. Results are presented as mean ± standard deviation (SD), with significance set at *p* < 0.05. Data normality was assessed using the Shapiro-Wilk test.

The analyses focused on repeated measures across six contraction levels: 5%, 10%, 20%, 30%, 50%, and 70% MVC. The effect of the contraction level was investigated for power spectral density, coherence, transmission delay, burst rate and duration, and related measures using Linear Mixed Models (LMM). For each dependant variable, a LMM was built with contraction level defined as fixed effect, and the subject as random effect. For all LMMs, the Fixed Effects Omnibus Test was used to evaluate the overall influence of contraction levels. When the omnibus test was significant (*p* < 0.05), post-hoc pairwise comparisons with Bonferroni correction for multiple comparisons were performed to identify specific differences between contraction levels. Post-hoc p-values are provided only for relevant comparisons where not all levels showed significant differences. For tests with *p* < 0.05, the corresponding F-values are reported. A linear regression model was used to evaluate the relationship between CMC and IMC. Additionally, a one-way ANOVA was conducted to evaluate significant differences in correlation coefficients between cortical ON bursting events and their corresponding peripheral delayed windows, non-delayed windows, and OFF windows where no bursting event was detected.

## Results

We extracted MU activity for each subject across different force levels, with the number of reliable MUs identified at each level reported in Table 1. To ensure consistency in neural drive estimation, we randomly selected a subset of units at each force level, matching the total number of action potentials analyzed across conditions (see Methods). This is needed to match the quality of estimate of the neural drive for each contraction level, which otherwise would be an influencing factor in coherence estimates (Negro et al., 2009) (Table 1).

**Table 1.** Summary of Experimental Results. Mean ± standard deviation of experimental results across six contraction levels, expressed as a percentage of maximum voluntary contraction (MVC). Power spectral density (PSD) values for electroencephalography (EEG) are scaled by 10^−3^, while those for the Cumulative Spike Train (CST) are scaled by 10^−4^.

|  | 5% MVC | 10% MVC | 20% MVC | 30% MVC | 50% MVC | 70% MVC |
| --- | --- | --- | --- | --- | --- | --- |
| Decomposed Motor Units | 23.58±6.41 | 28.41±10.65 | 28.41±10.55 | 26.75±10.74 | 17.33±9.59 | 7.83±5.02 |
| Included in the analysis Motor Units | 13.75±7.66 | 12.41±7.22 | 11.91±6.74 | 11.5±6.57 | 9.58±6.70 | 7.58±5.08 |
| Recruitment Thresholds [%] | 3.68±0.98 | 6.72±1.84 | 13.39±1.43 | 22.14±3.34 | 33.8±5.72 | 48.13±6.07 |
| Corticomuscular Coherence (CMC) | 0.08±0.02 | 0.09±0.03 | 0.08±0.02 | 0.07±0.03 | 0.07±0.02 | 0.08±0.02 |
| CMC transmission frequency [Hz] | 21.91±5.23 | 21.41±5.08 | 21.5±5.91 | 21.16±4.82 | 18.66±4.9 | 20.41±4.9 |
| Intramuscular Coherence (IMC) | 0.23±0.13 | 0.22±0.11 | 0.17±0.08 | 0.18±0.11 | 0.15±0.05 | 0.19±0.10 |
| IMC transmission frequency [Hz] | 20.83±5.23 | 29.91±4.33 | 18.83±4.36 | 18.83±5.11 | 20.50±5.35 | 19±3.74 |
| PSD CST [W/Hz] | 2.22±1.42 | 2.44±1.68 | 2.77±2.27 | 2.66±1.95 | 2.28±1.6 | 2.03±1.13 |
| PSD EEG [W/Hz] | 1.83±1.08 | 1.75±0.74 | 1.87±1.12 | 1.74±1.07 | 1.5±0.51 | 1.49±0.47 |
| Bursts Rate CST [Hz] | 3.55±0.76 | 3.61±0.83 | 3.72±0.5 | 3.73±0.92 | 3.53±0.74 | 3.32±0.65 |
| Bursts Duration CST [ms] | 59.46±10.36 | 58.69±10.74 | 54.97±5.63 | 52.26±4.75 | 64.11±9.67 | 70.28±10.1 |
| Bursts Rate EEG [Hz] | 3.18±1.22 | 3.09±1.27 | 3.22±1.26 | 3.12±0.9 | 3.21±1.19 | 3.27±0.89 |
| Burst Duration EEG [ms] | 54.20±8.35 | 57.52±7.72 | 56.55±7.37 | 53.01±4.88 | 54.80±10.2 | 54.23±6.71 |
| Transmission Delays [ms] | 25.99±3.24 | 26.16±3.36 | 24.17±3.54 | 22.77±3.61 | 21.49±3.16 | 22.28±2.29 |

MUs were then categorized into groups based on their recruitment thresholds, as shown in Figure 3. With increasing levels of MVC, the average recruitment thresholds of the subsets of identified MUs increased (*F* (5,55) = 303; *p* < 0.001), as expected (all *p* < 0.001). At each force level, indeed, the decomposition algorithm would tend to identify the MUAPs with highest energy that are usually associated to the MUs with highest thresholds (Farina and Holobar, 2016). According to the size principle (Henneman, 1957), the groups of MUs with different RTs corresponded to MNs of different sizes.

### Cortical Beta Oscillation are Uniformly Projected to the Motor Neuron Pool

We analysed CMC across groups of MNs with different recruitment thresholds. In all subjects, CMC values were statistically significant. CMC values remained constant across groups and conditions (Figure 2A; *F* (5,55) = 1.41; *p* = 0.23), indicating uniform frequency coupling between the cortex and MNs regardless of MN size. Additionally, no change was observed in the peak transmission frequency of CMC, further supporting the stability and uniformity of this transmission (*F* (5,55) = 0.63; *p* = 0.67). Similarly, IMC values were statistically significant across subjects, with no differences observed between MN groups (Figure 2B; *F* (5,55) = 2.08; *p* = 0.08) nor in its peak transmission frequency (*F* (5,55) = 0.56; *p* = 0.72). The similar behaviour of CMC and IMC for MN of different sizes was further confirmed the analysis of the linear correlation between CMC and IMC values in the beta band across subjects and MVC levels (Figure 3). A strong linear correlation was observed between CMC and IMC (*F* (12,59) = 11.5; *r*^*2*^ = 0.7; *p* < 0.001), supporting the hypothesis that MN beta power is mainly transmitted through cortical projections.

**Figure 2.**
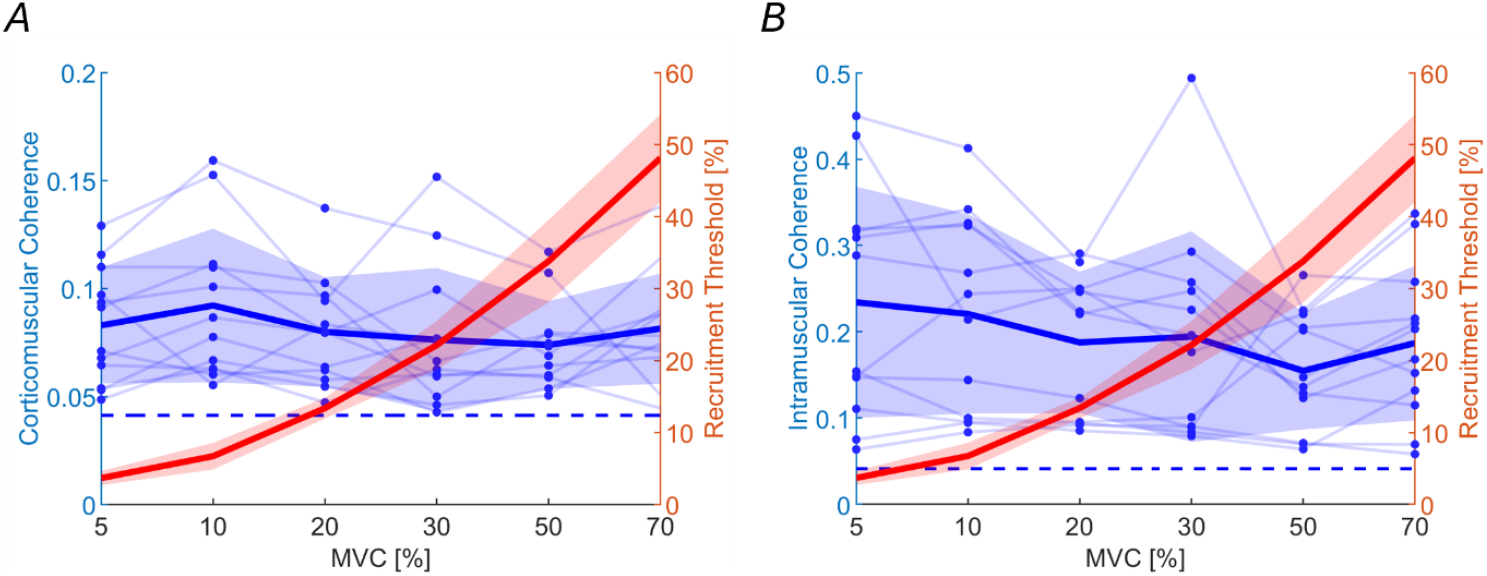
CMC and IMC do not change across different size Motor Units groups. Maximum corticomuscular coherence (CMC) and intramuscular coherence (IMC) values in the beta band range (13–30 Hz) are shown in blue, with individual data points and lines for each subject. Recruitment thresholds of the analysed motor units for each group are displayed in red. Bold lines indicate average values, while shaded areas represent the standard deviation. The horizontal dashed line marks the statistical significance threshold for the coherence values.

**Figure 3.**
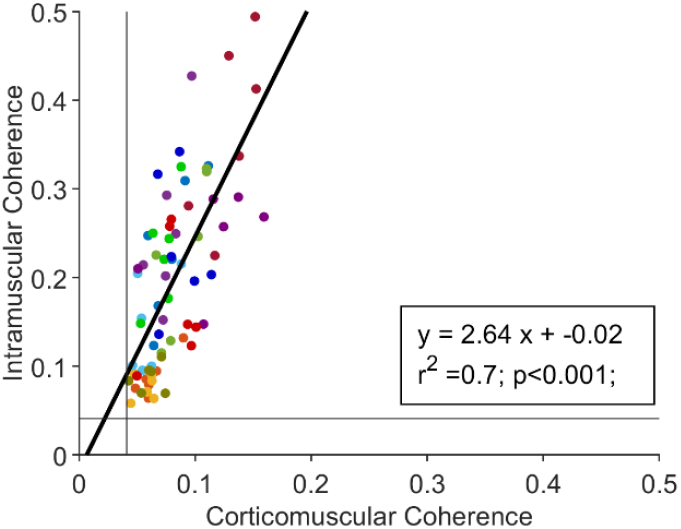
CMC and IMC values are linearly correlated. Each point represents the maximum intramuscular coherence (IMC) value in the beta band corresponding to the maximum corticomuscular coherence (CMC) value for each subject and contraction level. A total of 72 points are shown, color-coded by subject. The continuous light grey lines indicate the statistical significance threshold for coherence, which remains constant across conditions. The linear regression line is also displayed in black.

Because CMC is not directly related to the strength of the beta oscillations, being a normalised measure, we also analysed the power spectral density of EEG and the CST in the beta band across the different groups of MUs (Figure 4). The results showed no significant differences, with beta band power remaining stable in both the CST of the MU groups (Figure 4A, *F* (5,55) = 1.63; *p* = 0.17) and the EEG recorded from the ‘*Cz*’ channel (Figure 4B, *F* (5, 55) = 0.68; *p* = 0.64).

**Figure 4.**
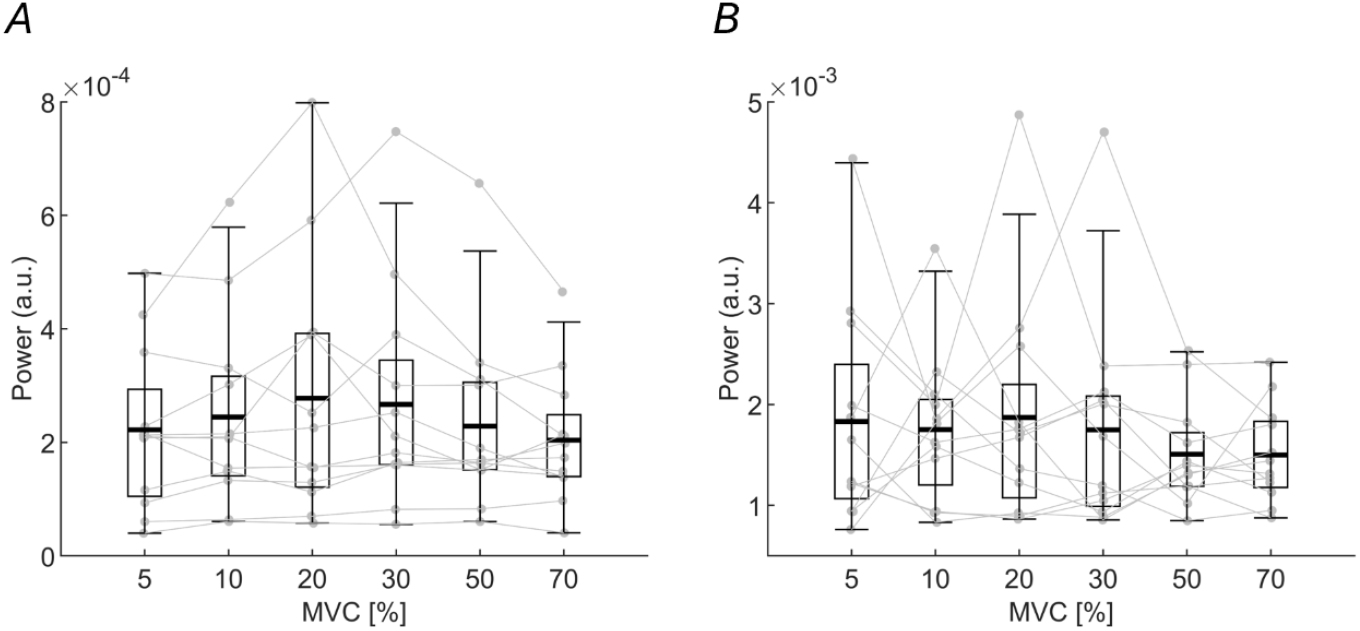
Power Spectral Density in the beta band across different contraction levels. Each point and line represent one subject. (A) Maximum values of the power spectral density in the beta band (13–30 Hz) across different contraction levels for the analysed groups of motor units. (B) The same analysis is made for the EEG. There are no significant changes.

During sustained contractions, cortical and peripheral activity occurs in bursts (Bräcklein et al., 2022; Echeverria-Altuna et al., 2022). Therefore, in addition to the spectral analysis described above, we also investigated the bursting-nature of central and peripheral beta oscillations. To determine whether the features of beta bursts, such as rate and duration, remained consistent across MNs of different sizes, we analyzed their stability. As shown in Figure 5 A,B, burst rates for both the CST and EEG remained stable across MN groups (*F* (5,55) = 0.97, *p* = 0.44; and *F* (5,55) = 0.06, *p* = 0.99; respectively). Burst duration also remained stable for EEG (Figure 5D; *F* (5,55) = 0.64; *p* = 0.67). However, a significant difference in CST burst duration was observed (Figure 5C; *F* (5,66) = 6.4; *p* < 0.001), in particular between 20% and 70% MVC (*p* = 0.001), and between 30% and 70% MVC levels (*p* < 0.001). All values for burst rate and duration are reported in Table 1.

**Figure 5.**
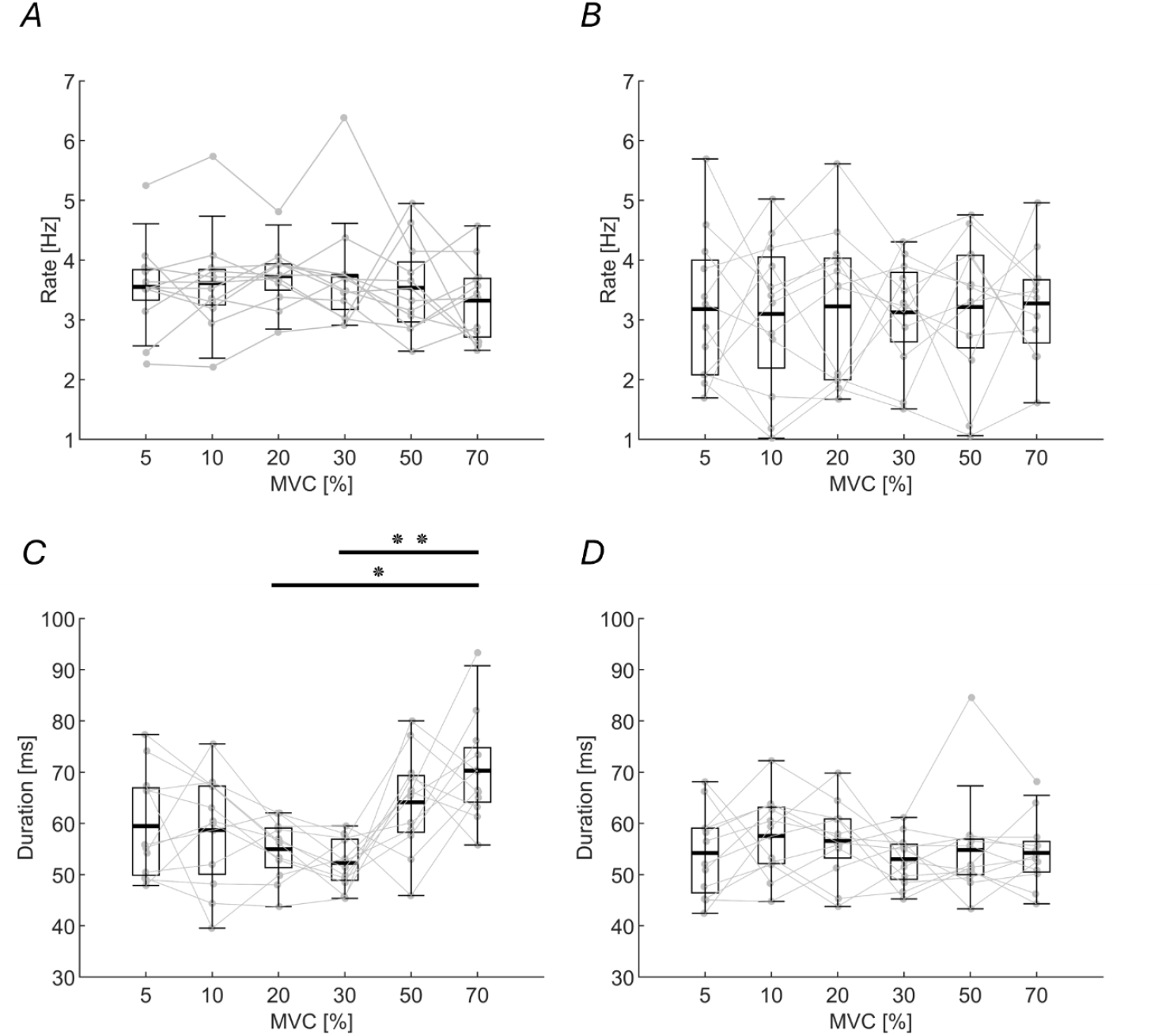
Beta burst rate and duration at cortical and peripheral level across different contraction levels. Each point and line represent an individual subject. This figure illustrates burst rate and duration values for motor unit (MU) groups and EEG recordings. (A) Average burst rate at the MU level. (B) Average burst rate at the ‘Cz’ EEG channel. (C) Average burst duration at the MU level, showing significant differences between 20% and 70% MVC, as well as between 30% and 70% MVC. (D) Average burst duration at the ‘Cz’ EEG channel. * *p* = 0.001; ** *p* < 0.001

### The Fastest Corticospinal Fibers Project Beta Oscillations to the entire Motor Neuron Pool

We further examined the transmission delay between the cortex and muscle. While previous findings showed that the fastest corticospinal fibers contribute to beta band CMC at low contraction levels (10% MVC) (Ibáñez et al., 2021), there are no previous results on beta transmission delays for higher threshold MNs. Here, we estimated the transmission delays with cumulant density analysis across contraction levels. The average cumulant density from the TA revealed a distinct positive peak in most individual traces at a positive time lag (Figure 6A). Significant cumulant density peaks were detected in all subjects and in 54 out of 72 analysed blocks.

**Figure 6.**
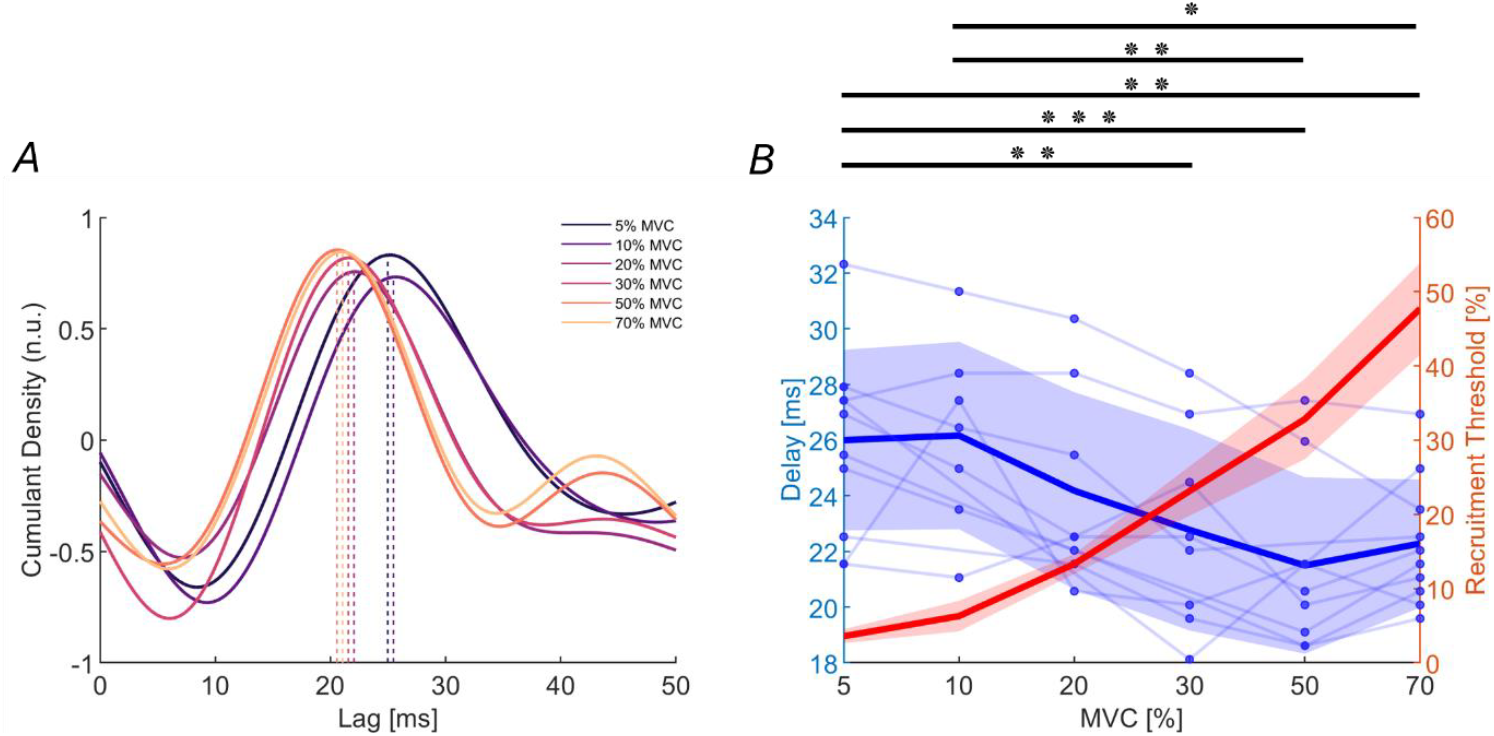
Transmission delays of cortical beta oscillations to the muscle show a significant decrease across Motor Units groups of different size. (A) Average normalized cumulant density across subjects for each contraction level between the EEG and the corresponding motor unit group. The dashed line represents the average peak and transmission delay identified using the cumulant density. (B) Transmission delays of beta band oscillations from the cortex to the tibialis anterior muscle, estimated using the cumulant density function, are shown in blue across different contraction levels, with individual subject data points and lines. The recruitment thresholds for the pooled motor units at each contraction level are shown in red. Solid lines indicate average values, while shaded areas represent the standard deviation. * *p* = 0.025; ** *p* < 0.01; *** *p* < 0.001.

The results revealed a significant reduction in transmission delay as the contraction intensity increased (Figure 6B; *F* (5,38.1) = 8.14; *p* < 0.001). The mean delays ranged from 26.0 ± 3.2 *ms* at 5% MVC to 22.3 ± 2.3 *ms* at 70% MVC, with intermediate values of 26.2 ± 3.4 *ms* (10% MVC), 24.2 ± 3.5 *ms* (20% MVC), 22.8 ± 3.6 *ms* (30% MVC), and 21.5 ± 3.2 *ms* (50% MVC). Significant differences in delay were observed between 5% and 30% MVC (*p* = 0.007); 5% and 50% MVC (*p* < 0.001); 5% and 70% MVC (*p* = 0.002); 10% and 50% MVC (*p* = 0.004); 10% and 70% MVC (*p* = 0.025) (Figure 6B). However, these differences in delay were fully compatible with the distribution of spinal MN axonal conduction velocities. Using an approximate average length of 500 *mm* for the axons of lower MNs innervating the TA and axonal conduction velocities ranging from 41 *m/s* at 5% MVC to 57.5 *m/s* at 70% MVC (Kakuda et al., 1992), the spinal MN axonal transmission delay varied between approximately 12.2 *ms* for lower-threshold MNs and 8.7 *ms* for higher threshold ones. This range of delays explains the range observed in estimates of corticomuscular delays, which resulted as the sum of a constant corticospinal delay of approximately 14 *ms* and a delay that depended on the variability in peripheral axonal conduction velocities (with the values reported above for low- and high-threshold MNs). This result supported the hypothesis that beta oscillations are transmitted homogenously to the entire MN pool using the fastest fibres of the corticospinal tract, regardless of MN size.

Finally, we extended our analysis to predict peripheral beta bursts appearance based on cortical bursts, accounting for the estimated average transmission delay between cortex and periphery. The reported burst rates for CST and EEG were comparable in number, supporting the idea that cortical bursts are transmitted to tonically contracted muscles, as previously reported (Bräcklein et al., 2022). For each subject, we identified cortical bursts and paired them with peripheral windows of equal duration, adjusted for the estimated average transmission delay at each contraction level (see Methods). We then calculated the correlation between these cortical bursts and their corresponding delayed ON windows, comparing them to OFF periods and non-delayed ON windows. Only significant correlation values were reported across subjects. Notably, 88.9% of blocks showed significant correlations for delayed ON windows, 79.1% for non-delayed ON windows, and 84.7% for OFF windows. Since we accounted for transmission delays at each MVC level and observed no significant differences in beta correlation or power between the cortex and TA across MVC levels (as shown in the results above), all averaged correlation coefficients were pooled across the three conditions. The median and standard deviation of correlation coefficients were 0.59 ± 0.56 for delayed ON windows, 0.17 ± 0.64 for non-delayed ON windows, and -0.40 ± 0.60 for OFF periods. The analysis revealed that correlation values for delayed ON windows were significantly greater than those for non-delayed ON windows (*p* = 0.039) and OFF periods (*p* < 0.001). A significant difference was also found between correlation coefficients of non-delayed ON windows and OFF periods (*p* = 0.043) (Figure 7). These findings further support that cortical beta bursts are transmitted to the periphery with delays that align with estimates from the cumulant density analysis.

**Figure 7.**
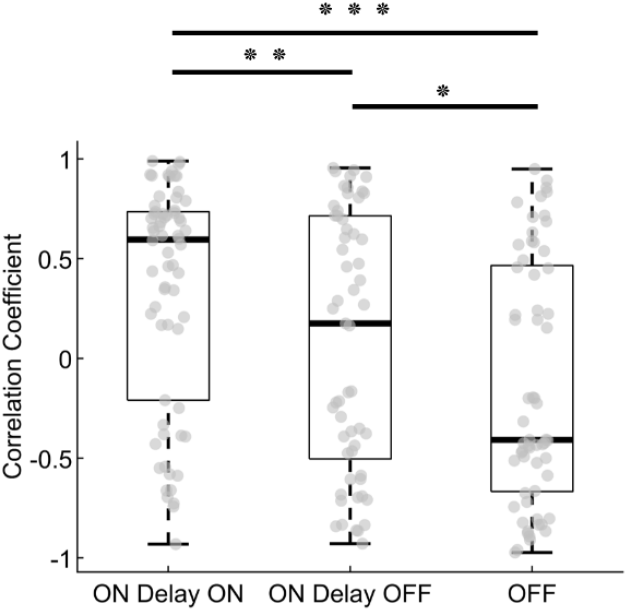
Correlation of cortical beta burst windows with peripheral delayed, non-delayed, and OFF windows. Significant correlation coefficients between cortical and corresponding peripheral burst windows for each contraction level and subject are shown in grey. Significant differences were found between cortical ON windows delayed by the average transmission delay across subjects for that contraction level (ON Delay ON) and non-delayed ON windows (ON Delay OFF), as well as between ON Delay ON windows and peripheral windows where no cortical burst occurred (OFF). * *p* = 0.043; ** *p* = 0.039; *** *p* < 0.001.

## Discussion

This study is the first to examine the corticospinal transmission of beta oscillations across distinct MN groups with markedly different recruitment thresholds, spanning nearly the entire force range of the muscle. Our findings demonstrate that cortical beta oscillations are transmitted uniformly across the MN pool, regardless of MN size. The estimated transmission delays align with the fastest corticospinal pathways. This uniform transmission remains consistent across different contraction levels, as confirmed by power spectral density analysis.

These results suggest that corticospinal fibers conveying beta oscillations engage all MNs similarly, utilizing the minimal delay enabled by the fastest pathways. Since CMC does not decrease with increasing recruitment threshold, it is likely that the corticospinal tract maintains its projection not only to low-threshold MUs but also to high-threshold ones. This finding challenges the assumption that high-threshold MNs—primarily recruited during strong contractions and largely controlled by the reticulospinal tract—receive less corticospinal input. Instead, our results indicate that beta band oscillations, like low-frequency common input involved in force control, are uniformly transmitted to the entire MN pool, reinforcing the widespread role of the corticospinal tract in motor control.

### Uniform Cortical Beta Oscillations Projection Across Different Motor Neuron Sizes

We identified a linear relationship between CMC and IMC values, suggesting that the primary source of peripheral beta oscillations is cortical activity. This finding aligns with previous studies demonstrating synchronization between cortical and peripheral beta oscillations (Bräcklein et al., 2022; Ibáñez et al., 2021). When the curve is populated with sufficient data from individual subjects, distinct clusters emerge, with each cluster corresponding to an individual. This observation supports earlier findings that CMC values vary between subjects (Matsuya et al., 2017), despite being unaffected by different MVC levels (see Results). Beyond confirming the cortical origin of peripheral beta oscillations, the linear CMC-IMC association implies that, with sufficient data, it may be possible to estimate CMC from IMC.

To accurately assess the corticomuscular transmission of beta oscillations across the MN pool, we accounted for and mitigated potential confounding factors in both cortical EEG signals and peripheral surface EMG signals — including phase correction to the former and motor unit decomposition to the latter. This approach enabled us to investigate whether beta band input to the MN pool is transmitted uniformly to spinal MNs, irrespective of their size, similar to the behaviour observed for low-frequency inputs linked to force control. Our findings revealed that CMC remained constant and independent of MN size. Furthermore, the power spectral density and bursting activity of beta oscillations were predominantly uniform across the MN pool.

### The Fastest Corticospinal Fibers Project to the Entire Motor Neuron Pool

Based on the measured neural dynamics, we concluded that cortical beta oscillations are uniformly projected across the MN pool. If this corticomuscular transmission is truly independent of MN size, it prompts further questions about the organization of upper MNs and their connectivity to the entire pool. To address this, we calculated the transmission delays of cortical beta oscillations across pools of MNs of varying sizes and observed a decrease in delay as MN size increased.

At low contraction levels, the transmission delays from cortex to muscle aligned closely with values previously reported using Motor Evoked Potentials (Cantone et al., 2019; Rossini et al., 1999). These delays are compatible only with the fastest corticospinal fibers transmitting cortical beta oscillations, consistent with prior findings (Ibáñez et al., 2021). However, these conclusions could previously be drawn only for low-threshold MUs.

Accounting for the peripheral axonal delays, which determine faster spinal-muscle transmission for larger MNs (see Results), our findings suggest that the corticospinal tract transmission delay remained constant. This consistent corticospinal transmission delay indicates that the same fastest corticospinal fibers projecting cortical beta oscillations to smaller MNs are also responsible for transmitting these oscillations to larger MNs. In summary, the central pathway delay is remarkably short, aligning with the conduction speed of the fastest corticospinal fibers, for both small and large MNs.

### Transmission of Cortical Beta Bursts to the Periphery

At the cortical level, beta burst rate and duration remained consistent across different contraction levels. However, while burst rate was constant across the MN pool, burst duration showed a significant increase between both 20% and 70% MVC, as well as 30% and 70% MVC. This is the only characteristic of beta oscillations and bursts that varied with MN size. While we cannot exclude physiological reasons, this variation can also be due to methodological limitations in estimating bursts durations in the two signals, with a situation of relatively poor signal-to-noise ratio.

We then analysed transmission of cortical bursts to the muscle. While Bräcklein et al. (2022) previously examined activity around cortical beta bursts, they did not include delay estimates. Here, we incorporated transmission delays, estimated using cumulant density, which varied with contraction levels. By applying a burst detection threshold to identify ON periods in cortical beta activity and estimating peripheral burst onsets using average transmission delays derived from the cumulant density analysis, we observed a stronger correlation between cortical and peripheral bursts compared to non-delayed ON periods and to OFF periods. This suggests that the transmission delays calculated here are likely accurate, as they identified time shifts that significantly increased the correlation between bursts at cortical and spinal levels. More importantly, this finding implies that the fastest corticospinal fibers projecting to the entire MN pool are also likely the primary pathway for cortical burst transmission. Nonetheless, the distribution of correlation values for ON periods delayed by the values identified with cumulant density still includes correlation coefficients near zero and even negative values. One plausible explanation is that not all cortical bursts identified as ON periods are effectively transmitted to the periphery. To address this limitation, future analyses could benefit from advanced burst detection algorithms capable of distinguishing bursts based on their waveform rather than a single amplitude threshold. Recent research has linked burst waveform characteristics at cortical level to specific functions in movement-related cortical dynamics (Szul et al., 2023), suggesting that analysing the specific waveforms of cortical bursts could help determine whether only a subset of them are transmitted to the periphery. Such approach could refine our understanding of the transmission dynamics between cortical and peripheral beta bursts.

### Limitations

The identification of heterogeneous and particularly large MUs remains a challenge due to the limitations of the EMG decomposition algorithm, particularly due to the increased cancellation of interference EMG signals at higher contraction levels. While larger MUs were successfully decomposed at submaximal contraction levels, even larger MUs likely remain to be identified at higher force levels — though such efforts are challenging both technically and due to muscle fatigue. Moreover, although the transmission delays at lower MVCs are consistent with values reported in the literature, it is important to note that the cumulant density function provides only a rough estimate of these delays, as it is sensitive to noise. Finally, all results were obtained for the TA muscle, and while beta band oscillations are ubiquitous, generalizing these findings to other muscles is not possible. Different muscles exhibit not only distinct anatomical features but also unique neural control strategies (Ushiyama et al., 2010). To determine the broader applicability of these conclusions, further studies using similar methods should be conducted across a variety of muscles.

## Acknowledgements

This work was supported by the European Research Council Synergy Grant Natural BionicS, Contract #810346. APV was supported by the European Union’s Horizon Europe research and innovation programme under the Marie Skłodowska-Curie grant agreement No 101151398. JIP was supported by the project ECHOES (European Research Council (ERC) Starting) under Grant 101077693 and by a Consolidación Investigadora project under Grant CNS2022-135366 funded by MCIN/AEI/10.13039/ 501100011033 and UE’s NextGenerationEU/PRTR Fu

